## Supplemental File 1 for "Intrinsic protein disorder is insufficient to drive subnuclear clustering in embryonic transcription factors": IDR_Identification_HMM.ipynb

```
# %% [code]
PATH = "/Users/eisen/software/iupred/"

# %% [code]
%matplotlib inline

from Bio import SeqIO
from Bio.Data import CodonTable
from Bio.SeqUtils import MeltingTemp as mt

import numpy as np
import scipy as sp

import subprocess
import os

import matplotlib

matplotlib.rcParams['text.usetex'] = False

import matplotlib.pyplot as plt
import math
import re

import sys
import textwrap
import math
import os

import pandas as pd

import seaborn as sns

from collections import Counter

import scipy.stats

import sklearn
import sklearn.preprocessing

import itertools 

import pickle

from pomegranate import *
from matplotlib.patches import Circle, Wedge, Polygon, Rectangle
from matplotlib.collections import PatchCollection

# %% [markdown]
#### Specify the directory containing your genome files:

# %% [code]
os.chdir("/Users/colleenhannon/Dropbox (Berkeley)/lcd")

# %% [code]
def avg(lst):
    return sum(lst) / len(lst)


def aa_freq(_seq):
    _freq = {}
    for _aa in _seq:
        if _aa in _freq:
            _freq[_aa] += 1
        else:
            _freq[_aa] = 1
    for _aa, _ins in _freq.items():
        _freq[_aa] = _ins / len(_seq)
    return _freq


def read_matrix(matrix_file):
    _mtx = {}
    with open(matrix_file, "r") as _fhm:
        for _line in _fhm:
            if _line.split()[0] in _mtx:
                _mtx[_line.split()[0]][_line.split()[1]] = float(_line.split()[2])
            else:
                _mtx[_line.split()[0]] = {}
                _mtx[_line.split()[0]][_line.split()[1]] = float(_line.split()[2])
    return _mtx


def read_histo(histo_file):
    hist = []
    h_min = float("inf")
    h_max = -float("inf")
    with open(histo_file, "r") as fnh:
        for _line in fnh:
            if _line.startswith("#"):
                continue
            if float(_line.split()[1]) < h_min:
                h_min = float(_line.split()[1])
            if float(_line.split()[1]) > h_max:
                h_max = float(_line.split()[1])
            hist.append(float(_line.split()[-1]))
    h_step = (h_max - h_min) / (len(hist))
    return hist, h_min, h_max, h_step


def smooth(energy_list, window):
    weighted_energy_score = [0] * len(energy_list)
    for idx in range(len(energy_list)):
        weighted_energy_score[idx] = avg(energy_list[max(0, idx - window):min(len(energy_list), idx + window + 1)])
    return weighted_energy_score

def read_seq(fasta_file):
    _seq = ""
    with open(fasta_file) as file_handler:
        for _line in file_handler:
            if _line.startswith(">"):
                continue
            _seq += _line.strip()
    return _seq

# %% [code]
def iupred(seq, mode):
    if mode == "short":
        lc = 1
        uc = 25
        wc = 10
        mtx = read_matrix("{}/data/iupred2_short_energy_matrix".format(PATH))
        histo, histo_min, histo_max, histo_step = read_histo("{}/data/short_histogram".format(PATH))

    elif mode == 'glob':
        lc = 1
        uc = 100
        wc = 15
        mtx = read_matrix("{}/data/iupred2_long_energy_matrix".format(PATH))
        histo, histo_min, histo_max, histo_step = read_histo("{}/data/long_histogram".format(PATH))

    else:
        lc = 1
        uc = 100
        wc = 10
        mtx = read_matrix("{}/data/iupred2_long_energy_matrix".format(PATH))
        histo, histo_min, histo_max, histo_step = read_histo("{}/data/long_histogram".format(PATH))

    unweighted_energy_score = [0] * len(seq)
    weighted_energy_score = [0] * len(seq)
    iupred_score = [0] * len(seq)

    for idx in range(len(seq)):
        freq_dct = aa_freq(seq[max(0, idx - uc):max(0, idx - lc)] + seq[idx + lc + 1:idx + uc + 1])
        for aa, freq in freq_dct.items():
            try:
                unweighted_energy_score[idx] += mtx[seq[idx]][aa] * freq
            except KeyError:
                unweighted_energy_score[idx] += 0

    if mode == 'short':
        for idx in range(len(seq)):
            for idx2 in range(idx - wc, idx + wc + 1):
                if idx2 < 0 or idx2 >= len(seq):
                    weighted_energy_score[idx] += -1.26
                else:
                    weighted_energy_score[idx] += unweighted_energy_score[idx2]
            weighted_energy_score[idx] /= len(range(idx - wc, idx + wc + 1))
    else:
        weighted_energy_score = smooth(unweighted_energy_score, wc)

    glob_text = ""
    if mode == 'glob':
        gr = []
        in_gr = False
        beg, end = 0, 0
        for idx, val in enumerate(weighted_energy_score):
            if in_gr and val <= 0.3:
                gr.append({0: beg, 1: end})
                in_gr = False
            elif in_gr:
                end += 1
            if val > 0.3 and not in_gr:
                beg = idx
                end = idx
                in_gr = True
        if in_gr:
            gr.append({0: beg, 1: end})
        mgr = []
        k = 0
        kk = k + 1
        if gr:
            beg = gr[0][0]
            end = gr[0][1]
        nr = len(gr)
        while k < nr:
            if kk < nr and gr[kk][0] - end < 45:
                beg = gr[k][0]
                end = gr[kk][1]
                kk += 1
            elif end - beg + 1 < 35:
                k += 1
                if k < nr:
                    beg = gr[k][0]
                    end = gr[k][1]
            else:
                mgr.append({0: beg, 1: end})
                k = kk
                kk += 1
                if k < nr:
                    beg = gr[k][0]
                    end = gr[k][1]
        seq = seq.lower()
        nr = 0
        res = ""
        for i in mgr:
            res += seq[nr:i[0]] + seq[i[0]:i[1] + 1].upper()
            nr = i[1] + 1
        res += seq[nr:]
        res = " ".join([res[i:i + 10] for i in range(0, len(res), 10)])
        glob_text += "Number of globular domains: {}\n".format(len(mgr))
        for n, i in enumerate(mgr):
            glob_text += "          globular domain   {}.\t{}-{}\n".format(n + 1, i[0] + 1, i[1] + 1)
        glob_text += "\n".join(textwrap.wrap(res, 70))

    for idx, val in enumerate(weighted_energy_score):
        if val <= histo_min + 2 * histo_step:
            iupred_score[idx] = 1
        elif val >= histo_max - 2 * histo_step:
            iupred_score[idx] = 0
        else:
            iupred_score[idx] = histo[int((weighted_energy_score[idx] - histo_min) * (1 / histo_step))]
    return iupred_score, glob_text


# %% [code]
def anchor2(seq, iupred_scores):
    local_window_size = 41
    iupred_window_size = 30
    local_smoothing_window = 5
    par_a = 0.0013
    par_b = 0.26
    par_c = 0.43
    iupred_limit = par_c - (par_a / par_b)
    mtx = read_matrix('{}/data/anchor2_energy_matrix'.format(PATH))
    interface_comp = {}
    with open('{}/data/anchor2_interface_comp'.format(PATH)) as _fn:
        for line in _fn:
            interface_comp[line.split()[1]] = float(line.split()[2])
    local_energy_score = [0] * len(seq)
    interface_energy_score = [0] * len(seq)
    energy_gain = [0] * len(seq)
    for idx in range(len(seq)):
        freq_dct = aa_freq(seq[max(0, idx - local_window_size):max(0, idx - 1)] + seq[idx + 2:idx + local_window_size + 1])
        for aa, freq in freq_dct.items():
            try:
                local_energy_score[idx] += mtx[seq[idx]][aa] * freq
            except KeyError:
                local_energy_score[idx] += 0
        for aa, freq in interface_comp.items():
            try:
                interface_energy_score[idx] += mtx[seq[idx]][aa] * freq
            except KeyError:
                interface_energy_score[idx] += 0
        energy_gain[idx] = local_energy_score[idx] - interface_energy_score[idx]
    iupred_scores = smooth(iupred_scores, iupred_window_size)
    energy_gain = smooth(smooth(energy_gain, local_smoothing_window), local_smoothing_window)
    anchor_score = [0] * len(seq)
    for idx in range(len(seq)):
        sign = 1
        if energy_gain[idx] < par_b and iupred_scores[idx] < par_c:
            sign = -1
        corr = 0
        if iupred_scores[idx] > iupred_limit and energy_gain[idx] < 0:
            corr = (par_a / (iupred_scores[idx] - par_c)) + par_b
        anchor_score[idx] = sign * (energy_gain[idx] + corr - par_b) * (iupred_scores[idx] - par_c)
        anchor_score[idx] = 1 / (1 + math.e ** (-22.97968 * (anchor_score[idx] - 0.0116)))
    return anchor_score


# %% [code]
genome_dir = "/Volumes/Data/Eisen Lab/flybase/"
genome_ver = "6.25"

# %% [code]
codontable ={
       'TTT': 'F', 'TTC': 'F', 'TTA': 'L', 'TTG': 'L', 
       'TCT': 'S', 'TCC': 'S', 'TCA': 'S', 'TCG': 'S', 
       'TAT': 'Y', 'TAC': 'Y', 'TAA': 'X', 'TAG': 'X',
       'TGT': 'C', 'TGC': 'C', 'TGA': 'X', 'TGG': 'W', 
       'CTT': 'L', 'CTC': 'L', 'CTA': 'L', 'CTG': 'L', 
       'CCT': 'P', 'CCC': 'P', 'CCA': 'P', 'CCG': 'P', 
       'CAT': 'H', 'CAC': 'H', 'CAA': 'Q', 'CAG': 'Q', 
       'CGT': 'R', 'CGC': 'R', 'CGA': 'R', 'CGG': 'R', 
       'ATT': 'I', 'ATC': 'I', 'ATA': 'I', 'ATG': 'M', 
       'ACT': 'T', 'ACC': 'T', 'ACA': 'T', 'ACG': 'T', 
       'AAT': 'N', 'AAC': 'N', 'AAA': 'K', 'AAG': 'K', 
       'AGT': 'S', 'AGC': 'S', 'AGA': 'R', 'AGG': 'R', 
       'GTT': 'V', 'GTC': 'V', 'GTA': 'V', 'GTG': 'V', 
       'GCT': 'A', 'GCC': 'A', 'GCA': 'A', 'GCG': 'A', 
       'GAT': 'D', 'GAC': 'D', 'GAA': 'E', 'GAG': 'E', 
       'GGT': 'G', 'GGC': 'G', 'GGA': 'G', 'GGG': 'G' 
}

codonlookup = {}

for aas in codontable.values():
    codonlookup[aas] = []
    
for codon in codontable:
    codonlookup[codontable[codon]].append(codon)
    

# %% [code]
aas = "ACDEFGHIKLMNPQRSTVWY"

# %% [code]
def reverse_complement(seq):
  """
  Return the reverse complement of the given DNA sequence.
  """
  T = {'A':'T', 'T':'A', 'G':'C', 'C':'G', 'N':'N'}
  s = ''
  for i in range(len(seq)-1, -1, -1):
    s += T[seq[i]]
  return s

# %% [code]
complement = {'A': 'T', 'C': 'G', 'G': 'C', 'T': 'A'} 

def reverse_complement(seq):    
    bases = list(seq) 
    bases = reversed([complement.get(base,base) for base in bases])
    bases = ''.join(bases)
    return bases

# %% [code]
class gff:
    
    def __init__(self):
        
        self.seq = ""
        self.source = ""
        self.feature = ""
        self.start = ""
        self.end = ""
        self.score = ""
        self.strand = ""
        self.frame = ""
        self.attribute = {}
    
    @classmethod
    def from_fields(cls,fields):
        #fields = line.split("\t")
        gff = cls()
        if len(fields) == 9:
            gff.seq = fields[0]
            gff.source = fields[1]
            gff.feature = fields[2]
            gff.start = int(fields[3])
            gff.end = int(fields[4])
            if fields[5] != ".":
                gff.score = float(fields[5])
            gff.strand = fields[6]
            gff.frame = fields[7]
            
            atts = fields[8].split(';')
            for att in atts:
                (k,v) = att.split('=')
                gff.attribute[k] = v.split(',')
        return gff

# %% [markdown]
## Parse current Drosophila gff file

# %% [code]
genome_dir = "/Volumes/Data/Eisen Lab/flybase/"
gff_file = "dmel-all-no-analysis-r6.25.gff"

gffs = {}

features = [
              'CDS',
              'enhancer',
              'exon',
              'five_prime_UTR',
              'gene',
              'golden_path_region',
              'insulator',
              'intergenic',
              'intron',
              'mRNA',
              'mature_peptide',
              'miRNA',
              'ncRNA',
              'none',
              'polyA_site',
              'pre_miRNA',
              'protein',
              'pseudogene',
              'rRNA',
              'regulatory_region',
              'repeat_region',
              'silencer',
              'snRNA',
              'snoRNA',
              'tRNA',
              'tandem_repeat',
              'three_prime_UTR',
              'transposable_element',
              ]

for feature in features:
    gffs[feature] = []

gffo = open(genome_dir + gff_file,"r")


for line in gffo:
    if line[0:2] != "##":
        line = line.strip()
        line = line.split("\t")
        if len(line) == 9:
            if line[2] in features:        
                gffs[line[2]].append(gff.from_fields(line))
                         

# %% [code]
gffs

# %% [code]
#
# parse current Drosophila gff file
#

genome_dir = "/Volumes/Data/Eisen Lab/flybase/"
gff_file = "dmel-all-filtered-r6.25.gff"

gffs_domains = {}

features = ['dmel_domain_PFAM','dmel_domain_SMART']

for feature in features:
    gffs_domains[feature] = []

gffo = open(genome_dir + gff_file,"r")

for line in gffo:
    if line[0:2] != "##":
        line = line.strip()
        line = line.split("\t")
        if len(line) == 9:
            if line[1] in features and line[2] == "match" :        
                gffs_domains[line[1]].append(gff.from_fields(line))
                         

# %% [code]
class protein:
    
    def __init__(self):
        
        self.id = ""
        self.name = ""
        self.parent = ""
        self.iep = ""
        self.mw = ""
        self.xref = {}
        self.seq = ""
        self.CDS = ""
        self.iupred = 0.0
        self.gff = ""

    @classmethod
    def from_atts(cls,atts):
        protein = cls()
        txt_fields = [('ID')]
        if 'ID' in atts:
            protein.id = atts['ID'][0]
        if 'Name' in atts:
            protein.name = atts['Name'][0]
        if 'Derives_from' in atts:
            protein.parent = atts['Derives_from'][0]
        if 'derived_isoelectric_point' in atts:
            protein.iep = float(atts['derived_isoelectric_point'][0])
        if 'derived_molecular_weight' in atts:
            protein.mw = float(atts['derived_molecular_weight'][0])
        if 'Dbxref' in atts:
            for xref in atts['Dbxref']:
                xrefs = xref.split(':')
                protein.xref[xrefs[0]] = xrefs[1]
    
        return protein
    
class mRNA:
    
    def __init__(self):
        
        self.id = ""
        self.name = ""
        self.parent = ""
        self.children = []
        self.score_text = ""
        self.xref = {}
        self.gff = ""
        self.seq = ""
        self.CDS = []
        
    @classmethod
    def from_atts(cls,atts):
        mRNA = cls()
        if 'ID' in atts:
            mRNA.id = atts['ID'][0]
        if 'Name' in atts:
            mRNA.name = atts['Name'][0]
        if 'Parent' in atts:
            mRNA.parent = atts['Parent'][0]
        if 'score_text' in atts:
            mRNA.score_text = atts['score_text'][0]
        if 'Dbxref' in atts:
            for xref in atts['Dbxref']:
                xrefs = xref.split(':')
                mRNA.xref[xref[0]] = xref[1]
    
        return mRNA

class gene:
    
    def __init__(self):
        self.id = ""
        self.name = ""
        self.fullname = ""
        self.aliases = []
        self.ontologies = []
        self.children = []
        self.score_text = ""
        self.xref = {}
        self.matzyg = ""
        self.stage2 = 0
        self.tf = 0
        self.proteins = []
        self.iupred = 0.0
        self.gff = ""
        
    @classmethod
    def from_atts(cls,atts):
        gene = cls()
        if 'ID' in atts:
            gene.id = atts['ID'][0]
        if 'Name' in atts:
            gene.name = atts['Name'][0]
        if 'score_text' in atts:
            gene.iep = float(atts['score_text'][0])
        if 'Dbxref' in atts:
            for xref in atts['Dbxref']:
                xrefs = xref.split('=')
                gene.xref[xref[0]] = xref[1]
        if 'Alias' in atts:
            gene.aliases = atts['Alias']
        if 'Ontology_term' in atts:
            gene.ontologies = atts['Ontology_term']
    
        return gene

# %% [code]
#
# process gff data for proteins, mRNAs and genes
#

proteins = {}

for protein_gff in gffs['protein']:
    p = protein.from_atts(protein_gff.attribute)
    p.gff = protein_gff
    proteins[p.id] = p

mRNAs = {}

for mrna_gff in gffs['mRNA']:
    p = mRNA.from_atts(mrna_gff.attribute)
    p.gff = mrna_gff
    mRNAs[p.id] = p
    
    
genes = {}

for gene_gff in gffs['gene']:
    p = gene.from_atts(gene_gff.attribute)
    p.gff = gene_gff
    genes[p.id] = p
    

# %% [code]
# set parents/children

for p in proteins:
    mRNAs[proteins[p].parent].children.append(proteins[p].id)
    
for t in mRNAs:
    genes[mRNAs[t].parent].children.append(mRNAs[t].id)
    
for g in genes.values():
    for m in g.children:
        for p in mRNAs[m].children:
            g.proteins.append(p)

# %% [code]
# set up hash to use attributes of genes or proteins to find gene

gene_lookup = {}
protein_lookup = {}

for g in genes.values():
    gene_lookup[g.id] = g.id
    gene_lookup[g.name] = g.id
    for a in g.aliases:
        gene_lookup[a] = g.id
    for x in g.xref.values():
        gene_lookup[x] = g.id
    for mid in g.children:
        m = mRNAs[mid]
        gene_lookup[m.id] = g.id
        gene_lookup[m.name] = g.id
        for x in m.xref.values():
            gene_lookup[x] = g.id
        for pid in m.children:
            p = proteins[pid]
            gene_lookup[p.id] = g.id
            protein_lookup[p.id] = p.id
            gene_lookup[p.name] = g.id
            protein_lookup[p.name] = p.id
            for x in p.xref.values():
                gene_lookup[x] = g.id
                protein_lookup[x] = p.id
        

# %% [code]
gene_lookup['FBtr0337064']

# %% [code]
len(gene_lookup.keys())

# %% [code]
for record in SeqIO.parse("/Volumes/Data/Eisen Lab/flybase/dmel-all-translation-r6.25.fasta", "fasta"):
    rs = record.description.split(" ")
    fbpp = rs[0]
    if fbpp in proteins:
        proteins[fbpp].seq = str(record.seq)

# %% [code]
cdsexons = {}

for record in SeqIO.parse("/Volumes/Data/Eisen Lab/flybase/dmel-all-CDS-r6.25.fasta", "fasta"):
    cdsexon = []
    fbpp == ""
    rs = record.description.split(" ")
    for r in rs:
        if r[0:6] == "dbxref":
            rr = r[7:].split(",")
            for r1 in rr:
                r1s = r1.split(":")
                if r1s[0] == "FlyBase":
                    fbpp = r1s[1]
                    if fbpp in proteins:
                        proteins[fbpp].CDS = str(record.seq)
        if r[0:3] == "loc":
            try:
                rr = r[4:-1].split(':')
                i = str(rr[1])

                c = rr[0]
                cdsexon.append(c)
                s = 1
                if i.startswith("complement"):
                    s = -1
                    i = i.replace("complement","")
                    i = i[1:-1]
                cdsexon.append(s)


                if i.startswith("join"):
                    i = i.replace("join","")
                    i = i[1:-1]

                    for si in i.split(","):
                        ssi = si.split("..")
                        cdsexon.append([int(x) for x in ssi])
                else:
                    ssi = i.split("..")
                    cdsexon.append([int(x) for x in ssi])
            except:
                print (rs)
                cdsexon = []
                
    if fbpp != "":
        cdsexons[fbpp] = [x for x in cdsexon]
    

# %% [code]
len(cdsexons.keys())

# %% [code]
j = []

i = "complement(join(19958500..19958900,19959301..19959367))"
i = "complement(20001103..20001966)"
s = 1
if i.startswith("complement"):
    s = -1
    i = i.replace("complement","")
    
print (s,i)
i = i[1:-1]
print (i)

if i.startswith("join"):
    i = i.replace("join","")
    i = i[1:-1]
    
    for si in i.split(","):
        ssi = si.split("..")
        j.append([int(x) for x in ssi])
else:
    print (i)
    ssi = i.split("..")
    j.append([int(x) for x in ssi])
        
    
print(j)

# %% [code]
genome_seq = {}
for record in SeqIO.parse("%s/dmel-all-chromosome-r%s.fasta" % (genome_dir,genome_ver), "fasta"):
    genome_seq[record.id] = str(record.seq)

# %% [code]
cds = {}
for cd_gff in gffs['CDS']:
    for cd_transcript in cd_gff.attribute['Parent']:
        if cd_transcript not in cds:
            cds[cd_transcript] = []
        cds[cd_transcript].append([cd_gff.start,cd_gff.end,cd_gff.strand])

# %% [code]
def cd_aa(cds,pos,strand):
    
    cds = sorted(cds,key=lambda x: x[0])
    
    t = np.sum([x[1] - x[0] + 1 for x in cds])
    
    cdpos = 0
    for x in cds:
        if pos >= x[0] and pos <= x[1]:
            cdpos += pos - x[0] + 1
            break
        else:
            cdpos += x[1] - x[0] + 1 
        
    if strand == "+":
        return int(cdpos/3)
    else:
        return int((t-cdpos)/3)

# %% [code]
domain_matches = {}

for p in proteins:
    domain_matches[p] = []

for feature in ['dmel_domain_PFAM','dmel_domain_SMART']:
    for gffline in gffs_domains[feature]:
        parents = gffline.attribute['Parent']
        dna = gffline.attribute['domain_name_abbreviation'][0]
        mid = gffline.attribute['ID'][0]
        for parent in parents:
            tx = proteins[parent].parent
            aa_pos = sorted([cd_aa(cds[tx],gffline.start,gffline.strand),cd_aa(cds[tx],gffline.end,gffline.strand)])
                    
            if parent not in domain_matches:
                domain_matches[parent] = []
            domain_matches[parent].append([aa_pos[0],aa_pos[1],feature,dna,mid,gffline.start,gffline.end])
            
indomain = {}

for p in proteins:
    indomain[p] = np.zeros(len(proteins[p].seq))
    
    for dm in domain_matches[p]:
        d_start = dm[0]
        d_end = dm[1]
        try:
            for x in range(d_start,d_end):
                indomain[p][x] = 1
        except:
            print (p,d_start,d_end)
        

# %% [code]
indomain

# %% [code]
len(indomain)

# %% [markdown]
## Compute IUPRED score for all proteins using three different IUPRED versions
### using local version of IUPRED -- see above

# %% [markdown]
#### Run IUPred long

# %% [code]
iupredout_long = {}

for g in genes:
    for p in genes[g].proteins:
        iupredout_long[p] = iupred(proteins[p].seq,'long')

 

# %% [markdown]
#### Write IUPRed output to .txt file

# %% [code]
f = open("iupredout_long.txt","w")
for p in proteins:
    d = iupredout_long[p][0]
    dd = " ".join([str(x) for x in d])
    f.write("%s\t%s\n" % (p,dd))
f.close()


# %% [markdown]
### If you have already run IUPred, you can load the output instead: 
#### Load the iupred_long.txt file

# %% [code]
# t = pd.read_csv('iupredout_long.txt',sep='\t',names=['protein','iupred'])

# iupredout_long = {}
# for i,r in t.iterrows():
#     iupredout_long[r.protein] = [[float(x) for x in r.iupred.split(' ')]]

# %% [code]
iupredout_long

# %% [code]
len(iupredout_long)

# %% [markdown]
#### Run IUPred Short

# %% [code]
iupredout_short = {}

for g in genes:
    for p in genes[g].proteins:
        iupredout_short[p] = iupred(proteins[p].seq,'short')


# %% [markdown]
#### Write the IUPred short output to a text file

# %% [code]
f = open("iupredout_short.txt","w")
for p in proteins:
    d = iupredout_short[p][0]
    dd = " ".join([str(x) for x in d])
    f.write("%s\t%s\n" % (p,dd))
f.close()


# %% [markdown]
#### Load the IUPred short output if you already have it:

# %% [code]
# t = pd.read_csv('iupredout_short.txt',sep='\t',names=['protein','iupred'])

# iupredout_short = {}
# for i,r in t.iterrows():
#     iupredout_short[r.protein] = [[float(x) for x in r.iupred.split(' ')]]

# %% [markdown]
#### Run IUPred glob

# %% [code]
iupredout_glob = {}

for g in genes:
    for p in genes[g].proteins:
        iupredout_glob[p] = iupred(proteins[p].seq,'glob')

# %% [markdown]
#### Write the IUPred glob output to a text file:

# %% [code]
f = open("iupredout_glob.txt","w")
for p in proteins:
    d = iupredout_glob[p][0]
    dd = " ".join([str(x) for x in d])
    f.write("%s\t%s\n" % (p,dd))
f.close()

# %% [markdown]
#### Load the IUPred glob output if you already have it:

# %% [code]
# t = pd.read_csv('iupredout_glob.txt',sep='\t',names=['protein','iupred'])

# iupredout_glob = {}
# for i,r in t.iterrows():
#     iupredout_glob[r.protein] = [[float(x) for x in r.iupred.split(' ')]]

# %% [markdown]
### Parse proteome into regions inside and outside of annotated domains

# %% [code]
def parse_domains(fullrange,domains):
    
    outdomains = []
    
    inout = {}
    inout[fullrange[0]] = 0
    inout[fullrange[1]] = 0
    
    for domain in domains:
        s = domain[0]
        e = domain[1]
        
        if s not in inout:
            inout[s] = 0
        inout[s] += 1
        
        if e not in inout:
            inout[e] = 0
        inout[e] -= 1
        
    endpoints = sorted(inout.keys())
    
    inoutsum = {}
    
    ios = 0
     
    for epi, ep in enumerate(endpoints):
        ios += inout[ep]
        
        if ios == 0:        
            inoutsum[ep] = 0
        else:
            inoutsum[ep] = 1
    
    domain_start = fullrange[0]
    domain_type = inoutsum[domain_start]
    
    for epi, ep in enumerate(endpoints):
        if inoutsum[ep] != domain_type:
            outdomains.append((domain_start,ep,domain_type))
            domain_start = ep
            domain_type = inoutsum[ep]
    
    if domain_start != fullrange[1]:
        outdomains.append((domain_start,fullrange[1],domain_type))
    
    return outdomains
            

# %% [code]
protein_domains = {}

for p in domain_matches:
    prolen = len(proteins[p].seq)
    ds = []
    for dm in domain_matches[p]:
        ds.append([dm[0],dm[1]])
    protein_domains[p] = parse_domains((1,prolen),ds)

# %% [code]
protein_domains

# %% [code]
protein_domains['FBpp0311265']

# %% [code]
# Zld protein entries = 'FBpp0300192', 'FBpp0074576', 'FBpp0074577', 'FBpp0311065', 'FBpp0311265'

# %% [code]
longest_proteins = []
longest_prot_lookup = {}

for g in genes:
    plist = []
    for p in genes[g].proteins:
        if "X" not in proteins[p].seq:
            plist.append(p)
    if len(plist) > 0:
        lp = plist[np.argmax([len(proteins[p].seq) for p in plist])]
        longest_proteins.append(lp)
        longest_prot_lookup[g] = lp

# %% [code]
len(longest_proteins)

# %% [markdown]
### Compute IUPRED scores in and outside of annotated domains

# %% [code]
iupreds = [[],[]]

for p in protein_domains:
    for d in protein_domains[p]:
        s = d[0] - 1 
        e = d[1] - 1
        t = d[2]
        
        for i in range(s,e):
            try:
                iupreds[t].append(iupredout_long[p][0][i])
            except:
                pass
    

# %% [code]
len(iupreds[0])

# %% [code]
iupreds

# %% [code]
len(protein_domains)

# %% [code]
#h = plt.hist(iupreds[0],bins=np.linspace(0,1,25),color='red',alpha=.25,density=True,label="not domains")
#h = plt.hist(iupreds[1],bins=np.linspace(0,1,25),color='green',alpha=.25,density=True,label="domains")
h = plt.hist(iupreds[0],bins=np.linspace(0,1,25),color='red',alpha=.25,density=True,label="Outside Domains")
h = plt.hist(iupreds[1],bins=np.linspace(0,1,25),color='green',alpha=.25,density=True,label="Inside Domains")
plt.legend()
#plt.title("IUPRED 'long' Scores in PFAM/SMART Domains")
plt.title("IUPred Scores in PFAM/SMART Domains")
plt.xlabel('IUPred Score')
plt.savefig("iupred_long_scores_histogram_new_labels.png", bbox_inches='tight', dpi=600)

# %% [code]
iupreds_short = [[],[]]

for p in protein_domains:
    for d in protein_domains[p]:
        s = d[0] - 1 
        e = d[1] - 1
        t = d[2]
        
        for i in range(s,e):
            try:
                iupreds_short[t].append(iupredout_short[p][0][i])
            except:
                pass


# %% [code]
h = plt.hist(iupreds_short[0],bins=np.linspace(0,1,25),color='red',alpha=.25,density=True,label="not domains")
h = plt.hist(iupreds_short[1],bins=np.linspace(0,1,25),color='green',alpha=.25,density=True,label="domains")
plt.legend()
plt.title("IUPRED 'short' Scores in PFAM/SMART Domains")
plt.xlabel('IUPRED Score')
plt.savefig("iupred_short_scores_histogram.png", bbox_inches='tight', dpi=600)


# %% [code]
iupreds_glob = [[],[]]

for p in protein_domains:
    for d in protein_domains[p]:
        s = d[0] - 1 
        e = d[1] - 1
        t = d[2]
        
        for i in range(s,e):
            try:
                iupreds_glob[t].append(iupredout_glob[p][0][i])
            except:
                pass

h = plt.hist(iupreds_glob[0],bins=np.linspace(0,1,25),color='red',alpha=.25,density=True,label="not domains")
h = plt.hist(iupreds_glob[1],bins=np.linspace(0,1,25),color='green',alpha=.25,density=True,label="domains")
plt.legend()
plt.xlabel('IUPRED Score')

plt.title("IUPRED 'glob' Scores in PFAM/SMART Domains")
plt.savefig("iupred_glob_scores_histogram.png", bbox_inches='tight', dpi=600)


# %% [code]
iupreds_glob[1]

# %% [markdown]
## Run IDR Prediction HMM 

# %% [code]
np.linspace(0,1,25)

# %% [markdown]
#### Define the HMM

# %% [code]
s0 = State(NormalDistribution(.5,.3), name="s0")
s1 = State(NormalDistribution(.8,.3), name="s1")

model = HiddenMarkovModel('disorder iupred long')
model.add_states([s0, s1])
model.add_transition(model.start, s0, .5)
model.add_transition(model.start, s1, .5)
model.add_transition(s0, s0, .99)
model.add_transition(s0, s1, .01)
model.add_transition(s1, s0, .01)
model.add_transition(s1, s1, .99)

model.freeze()
model.bake()

# %% [markdown]
#### Calcuate Viterbi path

# %% [code]
iupred_vit_long = {}

for p in proteins:
    disorder_viterbi = model.viterbi(iupredout_long[p][0])
    disorder_viterbi_states = [i for i, state in disorder_viterbi[1]]
    disorder_viterbi_states.pop(0)
    iupred_vit_long[p] = disorder_viterbi_states
    

# %% [code]
s0 = State(NormalDistribution(.4,.3), name="s0")
s1 = State(NormalDistribution(.7,.3), name="s1")

model_short = HiddenMarkovModel('disorder iupred short')
model_short.add_states([s0, s1])
model_short.add_transition(model_short.start, s0, .5)
model_short.add_transition(model_short.start, s1, .5)
model_short.add_transition(s0, s0, .99)
model_short.add_transition(s0, s1, .01)
model_short.add_transition(s1, s0, .01)
model_short.add_transition(s1, s1, .99)

model_short.freeze()
model_short.bake()

# %% [code]
iupred_vit_short = {}

for p in proteins:
    disorder_viterbi = model_short.viterbi(iupredout_short[p][0])
    disorder_viterbi_states = [i for i, state in disorder_viterbi[1]]
    disorder_viterbi_states.pop(0)
    iupred_vit_short[p] = disorder_viterbi_states
    

# %% [code]
iup_0 = []
iup_1 = []

for p in proteins:
    iup_0.extend([i for i,j in zip(iupredout_long[p][0],iupred_vit_long[p]) if j == 0])
    iup_1.extend([i for i,j in zip(iupredout_long[p][0],iupred_vit_long[p]) if j == 1])

plt.hist(iup_0,alpha=.5)
plt.hist(iup_1,alpha=.5)

# %% [markdown]
### Plot HMM outputs

# %% [code]
def vit_plot(p):
    
    fig, ax = plt.subplots()
    
    ax.plot(iupredout_long[p][0], c= 'black',label='IUPred', alpha=.4) #long
    #ax.plot(iupredout_short[p][0], c='blue',label='IUPRED short', alpha=.4)
    #ax.plot(iupredout_glob[p][0], c='green',label='IUPRED glob', alpha=.4)
    ax.plot(iupred_vit_long[p], c='red',label='Viterbi') #long
    #ax.plot(iupred_vit_short[p],c='black',label='viterbi short')

    rects = []

    if p in protein_domains:
        for prd in protein_domains[p]:
            if prd[2] == 1:
                rects.append(Rectangle((prd[0],1.05),prd[1]-prd[0],.05))


    pc = PatchCollection(rects, facecolor='green', alpha=.25,edgecolor='black')
    ax.add_collection(pc)

    ax.set_ylim(0,1.1)
    
    plt.legend()
    ax.legend(loc='center left', bbox_to_anchor=(1, 0.5))

# %% [code]
vit_plot('FBpp0074577')

plt.xlabel('aa position')
plt.ylabel('IUPred score')

plt.savefig("FBpp0074577_Zld_vit_plot_final.png", bbox_inches='tight', dpi=600)


# %% [code]
iupreds = [[],[]]

for p in protein_domains:
    seq = proteins[p].seq
    for d in protein_domains[p]:
        s = d[0] - 1 
        e = d[1] - 1
        t = d[2]
        
        for i in range(s,e):
            try:
                iupreds[t].append(iupred_vit_long[p][i])
            except:
                pass
            
dom_vit = [Counter(iupreds[0]),Counter(iupreds[1])]

fig,ax = plt.subplots()

ax.bar([1,4],[dom_vit[1][0],dom_vit[0][0]],color='blue',label='Viterbi Structured')
ax.bar([2,5],[dom_vit[1][1],dom_vit[0][1]],color='red',label='Viterbi Unstructured')
ax.set_xticks([1.5,4.5])
ax.set_xticklabels(['PFAM/SMART Domains','Rest of Proteome'])
ax.set_title("Presence of Amino Acid in Domain vs. Viterbi Call")
ax.legend()
plt.savefig("iupred_long_viterbi_barplot.png", bbox_inches='tight', dpi=600)

# %% [code]
iupreds = [[],[]]

for p in protein_domains:
    seq = proteins[p].seq
    for d in protein_domains[p]:
        s = d[0] - 1 
        e = d[1] - 1
        t = d[2]
        
        for i in range(s,e):
            try:
                iupreds[t].append(iupred_vit_short[p][i])
            except:
                pass

dom_vit = [Counter(iupreds[0]),Counter(iupreds[1])]

fig,ax = plt.subplots()

ax.bar([1,4],[dom_vit[1][0],dom_vit[0][0]],color='blue',label='viterbi structured')
ax.bar([2,5],[dom_vit[1][1],dom_vit[0][1]],color='red',label='viterbi unstructured')
ax.set_xticks([1.5,4.5])
ax.set_xticklabels(['PFAM/SMART','Rest of proteome'])
ax.set_title("Presence of Amino Acid in Domain vs. Viterbi Call (short)")
ax.legend()
plt.savefig("iupred_short_viterbi_barplot.png", bbox_inches='tight', dpi=600)

# %% [markdown]
## Generate a list of IDRs for cloning

# %% [code]
domain_iupred = {}
domain_lcannot = {}
domain_seqs = {}
domain_info = {}


for feature in ['dmel_domain_PFAM','dmel_domain_SMART']:
    for gffline in gffs_domains[feature]:
        parents = gffline.attribute['Parent']
        dna = gffline.attribute['domain_name_abbreviation'][0]
        mid = gffline.attribute['ID'][0]
        for parent in parents:
            tx = proteins[parent].parent
            aa_pos = sorted([cd_aa(cds[tx],gffline.start,gffline.strand),cd_aa(cds[tx],gffline.end,gffline.strand)])
                    
            if parent in longest_proteins:
                if dna not in domain_iupred:
                    domain_iupred[dna] = []
                    domain_lcannot[dna] = []
                    domain_seqs[dna] = []
                    domain_info[dna] = []
                    
                
                d_start,d_end = [int(x)-1 for x in aa_pos]

                for i in range(d_start,d_end):
                    domain_iupred[dna].append(iupredout_long[parent][0][i])
                    domain_lcannot[dna].append(iupred_vit_long[parent][i])

                seq = proteins[parent].seq[d_start:d_end]
                domain_seqs[dna].append(seq)
                domain_info[dna].append([proteins[parent].parent,parent,aa_pos,len(seq)])


# %% [code]
ddata = []

for d in domain_lcannot:
    if len(domain_lcannot[d]) > 100:
        ddata.append([d,len(domain_seqs[d]),len(domain_lcannot[d]),np.average(domain_lcannot[d]),np.average(domain_iupred[d])])
        
domain_df = pd.DataFrame(ddata,columns=['domain','n','aas','fdisorder','iupred'])        
domain_df = domain_df.set_index('domain')

# %% [code]


# %% [code]
iupred_vit_domains = [[],[]]

for p in longest_proteins:
    pos = 0
    for k,g in itertools.groupby(iupred_vit_long[p]):
        gl = list(g)
        glen = len(gl)
        s = pos
        e = pos + glen
        seq = proteins[p].seq[s:e]
        iupred_vit_domains[k].append([p,len(proteins[p].seq),s,e,np.average(iupredout_long[p][0][s:e]),np.average(indomain[p][s:e]),seq])
        pos += glen
        

# %% [code]
ivdomains = []

for p in longest_proteins:
    pos = 0
    for k,g in itertools.groupby(iupred_vit_long[p]):
        gl = list(g)
        glen = len(gl)
        s = pos
        e = pos + glen
        seq = proteins[p].seq[s:e]
        ivdomains.append([k,p,len(proteins[p].seq),glen,s,e-1,np.average(iupredout_long[p][0][s:e]),np.average(indomain[p][s:e]),seq])
        pos += glen
        
ivdomains = pd.DataFrame(data = ivdomains, columns=['type','protein','prolen','domlen','domainstart','domainend','iupred','indomain','seq'])

# %% [code]
ivdomains.head(20)

# %% [code]
# Filter proteins by TFs
flytfs = pd.read_csv("flytfs.txt")

# %% [code]
fbpps = []
lps = []

for cg in flytfs.CG.values:
    fbpps.append(gene_lookup[cg])
    if gene_lookup[cg] in longest_prot_lookup:
        lps.append(longest_prot_lookup[gene_lookup[cg]])
    else:
        lps.append("")
        print (gene_lookup[cg])
flytfs['gene'] = fbpps
flytfs['lprot'] = lps

# %% [code]
tfprots = flytfs[flytfs.lprot != ""].lprot.values

# %% [code]


# %% [code]
inds = []

for i,r in ivdomains.iterrows():
    if r.protein in tfprots:
        inds.append(i)
        
ivdomains['tf'] = 0
ivdomains.loc[inds,'tf'] = 1
        

# %% [code]
# This is just to get a list of the PFAM domains and IDRs together.
tf_domains = ivdomains[(ivdomains.tf == 1)].copy()

# %% [code]
tf_domains

# %% [code]
tf_domains[tf_domains.protein == 'FBpp0084158']

# %% [code]
lcds_toclone = ivdomains[(ivdomains.type == 1) & (ivdomains.tf == 1) & (ivdomains.domlen >= 200) & (ivdomains.domlen <=400)].copy()

# %% [code]
len(lcds_toclone)

# %% [code]
inds = []
seqs = []

lcds_toclone['dnaseq'] = ""

for i,r in lcds_toclone.iterrows():
    s = r.domainstart
    e = r.domainend
    sn = 3 * s
    en = 3 * (e+1)
    if proteins[r.protein].CDS != "":
        try:
            ps = proteins[r.protein].CDS
            ns = ps[sn:en]
            inds.append(i)
            seqs.append(ns)
        except:
            print (r.protein)


# %% [code]
lcds_toclone.loc[inds,'dnaseq'] = seqs

# %% [code]
lcds_toclone.head(10)

# %% [code]
i = 4

r = lcds_toclone.iloc[i]

print (r.dnaseq)

# %% [code]
from Bio.Seq import Seq
from Bio.Alphabet import IUPAC

for i,r in lcds_toclone.iterrows():
    dna = r.dnaseq
    dna_bp = Seq(dna, IUPAC.unambiguous_dna)
    tdna = str(dna_bp.translate())
    
    if r.seq != tdna:
        print (r.protein,r.seq,tdna)

# %% [code]
len(lcds_toclone)

# %% [code]
inds = []

for i,r in lcds_toclone.iterrows():
    d = r.dnaseq
    g = 0
    for c in ['2L', '2R', '3L', '3R', '4', 'X']:
        if d in genome_seq[c]:
            g = 1
            break
        if reverse_complement(d) in genome_seq[c]:
            g = 1
            break
    if g == 1:
        inds.append(i)
    
lcds_toclone['genomic'] = 0
lcds_toclone.loc[inds,'genomic'] = 1

# %% [code]
lcds_toclone[lcds_toclone.genomic == 1]

# %% [code]
lcds_toclone.to_csv("lcds_toclone.tsv",sep = "\t")

# %% [code]
```
